# Head engagement during visuomotor tracking is determined by postural demands and aging

**DOI:** 10.1101/2025.02.19.639184

**Authors:** Petros Georgiadis, Vassilia Hatzitaki, Katja Fiehler, Dimitris Voudouris

## Abstract

Vision is important for various tasks, from visually tracking moving objects to maintaining balance. People obtain visual information through eye movements performed either alone or in combination with head movements. Even when isolated eye movements can accommodate the amplitude of the desired gaze shift, humans still perform head movements, as they provide additional sensory signals that can be integrated with retinal input resulting in improved gaze estimates. However, head movements also create mechanical torques and attenuate vestibular processing that could disturb balance. We, therefore, here examined whether head engagement is determined by postural requirements when performing a visual tracking task. Young participants visually tracked a target moving horizontally along different amplitudes, while they were seated, standing on a firm and an unstable surface. Our results showed stronger head engagement when standing than sitting, but no systematic differences were found between firm and unstable surfaces. To further explore the interplay between head engagement and postural demands, we conducted a second experiment where young and older participants performed a similar task, but now they were either allowed to move their head or instructed to limit their head movements. Both tracking accuracy and postural sway increased when engaging the head. When asked to limit head movements, both age groups engaged their head minimally, but head movements were more pronounced in more challenging postures. When allowed to move their head naturally, younger participants engaged their head more when standing than sitting, but older adults reduced their head movements with more demanding postures. We suggest that head movements in younger adults facilitate visual tracking, while limited head movements in older adults preserve balance.

## Introduction

Vision dominates our actions, such as eye (Land & Tatler, 2009), hand (Karl & Whishaw, 2013; Voudouris et al., 2018), or full-body movements (Mann et al., 2013). Humans typically obtain visual input by combining eye and head movements that shift gaze to positions of interest. Although gaze shifts are important to bring relevant information within the high-resolution fovea, visual information processing during gaze shifts is compromised, for instance, due to the fast eye movements blurring the visual image. Moreover, when visually tracking a moving target with the head restrained, the background moves across the retina in the opposite direction of the target’s motion, generating a relative motion signal that impedes visual perception and spatial orientation (Kowler, 2011). Such eye movements can have various implications for sensorimotor control. For instance, visually pursuing a moving target while standing with the head fixed leads to greater center of pressure (CoP) displacements both in healthy young and older adults (Thomas et al., 2017; Thomas et al., 2016). However, humans rarely shift their gaze with their head restrained. Although relatively small gaze shifts of about 15 degrees are typically accommodated by isolated eye movements (Hallet, 1986), larger gaze shifts are performed with combined eye-head movements (Daye et al., 2012; Pallus & Freedman, 2016; Georgiadis et al., 2023), even when the desired gaze shift can still be performed without these head movements.

Rotating the head when shifting gaze can have implications for postural control. On the one hand, head rotations provide neck proprioception signals that can be integrated with retinal and extra-retinal signals and lead to more reliable estimates of gaze orientation (Saglam et al., 2011). It is, thus, not surprising that humans visually track a moving baseball by employing head rotations and that s are more pronounced in expert players (Mann et al., 2013). Thus, head rotations may facilitate tasks that rely on visual information, such as keeping an upright stance while visually tracking a moving object (Raffi & Piras, 2019). On the other hand, as the head makes up ∼8% of total body mass (Winter, 2009), head rotations can create mechanical torques that could disturb body mechanics and increase postural sway (Fukushima et al., 2008). Meanwhile, voluntary movements can suppress self-generated sensory information (Cullen 2019; Blakemore et al., 2000). In the case of head rotations, the neck muscles’ motor command to rotate the head is integrated with retinal input and can suppress vestibular activation (Cullen, 2019). This vestibular suppression due to head rotations for a gaze shift could degrade spatial body representations and compromise body balance (Lopez et al., 2015). Given the complex interaction between the visual, vestibular, and proprioceptive systems associated with postural control (Peterka, 2018), one can only speculate on whether postural demands influence the engagement of the head when performing visuomotor tasks.

There is some indirect evidence that postural demands may alter how humans employ their head when performing visuomotor tasks. For example, head engagement is stronger when walking through more complex terrains (Thomas et al., 2020) which presumably increases postural challenges. More specifically, individuals demonstrate increased downward head orientations when walking on more challenging (i.e. rough, uneven) terrains, but this is likely due to more complex terrains requiring accurate foot placement and thus a clearer visual representation of the upcoming foothold (Matthis et al., 2018). In these cases, humans need to direct their gaze to the ground just in front of them, which can only be achieved by large downward head rotations. Increased postural demands arise also during healthy aging (Wang et al., 2024), which might explain why older adults rotate their head more when shifting gaze during walking (Cinelli et al., 2008; Hirasaki et al., 1993), and when visually tracking a moving target while standing (Georgiadis et al., 2023). However, it is not clear how head engagement is determined by postural demands.

To examine the influence of postural demands on head engagement, in our first experiment, young adults pursued a visual target moving along three different horizontal amplitudes under three distinct postural demands: sitting, standing on a firm surface, and standing on an unstable surface (foam). In line with previous results, we expected larger head rotations with increased amplitudes of the moving target (Freedman, 2008) and larger CoP displacements when standing on an unstable rather than a firm surface (Patel et al., 2008). If increased postural challenges require smaller head rotations, to reduce possible postural disturbances, for instance, we would expect reduced head movements when standing than sitting, and when standing on unstable than firm surfaces. If postural imbalance calls for stronger head engagement, for instance, to improve gaze estimates, we expect larger head movements when standing than sitting and when standing on unstable than firm surfaces.

To further investigate the potential consequences of the head’s engagement in visual tracking and postural control, we conducted a second experiment, where participants tracked a moving target either with their head allowed to move naturally (as in Experiment 1) or with the instruction to keep their head fixed. This set-up was selected to explicitly explore the role of adjusting head engagement in more demanding postures and its implications. If head engagement during tracking directly relates to CoP sway, we should observe smaller CoP sway when tracking with the head fixed. We now also asked both young and older adults to perform these two tracking tasks. We did so because healthy aging naturally increases postural instability as sensorimotor processes decline with age (Wang et al., 2024). As a result, older adults prioritize posture over other sensorimotor or cognitive tasks that are concurrently performed (Brown et al., 2002; Doumas et al., 2008). For example, older adults rotate their head more when shifting gaze during standing or walking compared to younger individuals (Cinelli et al., 2008; Hirasaki et al., 1993). However, to the best of our knowledge, there are no studies investigating how postural demands influence head engagement strategies during visual tracking tasks in aging.

In this second experiment, therefore, we examined the interplay between head engagement and balance control, as well as the role of aging in modulating the relationship between the two. We used the same experimental setup as in Experiment 1. A new group of young adults and a group of healthy older adults participated in the experiment. We included two conditions concerning the engagement of the head: a) head-free, where participants were allowed to naturally engage their heads without any restrictions, as in Experiment 1, and b) head-fixed, where participants were instructed to restrain head movements. If head engagement in the visual tracking task leads to greater postural sway, we expect greater CoP sway in the head-free compared to the head-fixed condition. In addition, based on previous findings (Cinelli et al., 2008; Hirasaki et al., 1993), we expected distinct age-related differences in head engagement strategies during tracking. If head engagement in the tracking task increases CoP sway, we expect older adults, who typically prioritize postural control (Brown et al., 2002; Doumas et al., 2008), to actively reduce their head engagement when performing under more demanding postures. Finally, if the engagement of the head holds apparent benefits for visual tracking performance, for instance, by providing more reliable gaze estimates (Saglam et al., 2011), we expect greater coupling between the gaze and target movements in the head-free than head-fixed condition.

## Methods

### Participants

The sample size was estimated a priori based on the expected effect of postural demands on the head’s engagement during a visual tracking task. After collecting pilot data of 5 young participants, who stood on a firm and on an unstable surface while tracking a target moving along three horizontal amplitudes (15, 42, and 95 degrees of visual angle), we considered a difference in the head engagement between the two postural conditions of 2.38 ± 0.49 degrees (mean ± SD). Combined with an α = 0.05 to achieve power = 0.90, a minimum of 24 participants (effect size = 0.96) was required. The power analysis was conducted in G*Power 3 (Faul et al., 2007). For experiment 1, we recruited a total of 28 healthy participants through student pools of Justus Liebig University Giessen. Due to high-frequency noise contaminating gaze and kinematic data, 4 participants were excluded from the analyses, resulting in a final sample size of 24 participants (6 males, mean age: 25 years ± 4.5, range 21-35). For experiment 2, a new set of eighteen younger (5 males, mean age: 24.3 years ± 3.3, range 20-33) and eighteen older adults (9 males, mean age: 63.1 years ± 5.4, range 55-71) were recruited. Young adults were, once again, obtained by the university student pools of Justus Liebig University Giessen, while none of them had joined Experiment 1. Older adults, who were community-dwelling, were recruited from internal mailing lists and personal contacts of the authors. Older adults were tested for cognitive functioning using the Montreal Cognitive Assessment, which they all completed with a score > 26 (Nasreddine et al., 2005), indicating normal cognitive ability. All participants had normal or corrected-to-normal vision without any known neurological or motor disorders that would impair their participation. Participants provided written consent and were compensated for their time, receiving either 8 Euros/hour (young and older participants) or course credits (only younger participants). The experiment was conducted in compliance with the guidelines of the local ethic’s committee of Justus Liebig University Giessen and the Declaration of Helsinki (2013; except for §35, pre-registration).

### Apparatus, stimuli, and protocol

For both experiments, participants were positioned inside a large room, facing a wall at 1.5 m. A projector (LG CineBeam LED Projector RGB LED, 1280 x 720 pixels, Screen Size: 25" ∼ 100", refresh rate: 60Hz) was placed about 3.5 m behind the participants and projected the visual stimulus (a red circle) on the wall that participants were facing. The room was darkened, with the lights turned off and the window blinds closed, which fostered the seamless blend of the projected screen into the dark surroundings, eliminating visible edges. However, some residual light was present in the room. The height of the stimulus presentation was adjusted to 10 degrees below the line of sight, depending on the participant’s height and the postural condition (seated, firm, foam; see below). Participants wore a portable eye-tracker (Tobii Pro Glasses 2) that recorded eye movements at 100 Hz, which was also equipped with two rigid bodies of three reflective markers each. An 8-camera system (Vicon Motion Systems, Oxford, UK) recorded at 100 Hz the position of these six reflective markers along with another two single markers that were attached to the right and left acromia of the participant. A force platform (AMTI, Massachusetts, US) recorded the vertical ground reaction force and moments in the mediolateral (ML) and the anteroposterior (AP) direction at 100 Hz to assess the CoP displacement. The design and presentation of visual stimuli was done with Psychtoolbox (Kleiner et al., 2007; version 3.0.18). The Vicon’s Software Development Kit (SDK) was used to establish communication between the Vicon system software (Nexus version 2.25.0) and Matlab R2023b (The Mathworks Inc., Massachusetts, US). Integrating all systems through Matlab allowed us to synchronize visual stimulus presentation with the sampling and digitization of the force, marker kinematics, and target motion signals via the Vicon’s data acquisition board (MX Giganet).

In experiment 1, young participants performed three counterbalanced blocks of 15 trials each, while they were at one of three postural configurations: they could either be seated on a chair, standing on a firm surface (directly on the force plate), or standing on an unstable surface (foam pad on the force plate; 50 x 40 x 6 cm; SISSEL BalanceFit Pad, novacare GmbH, Germany). Participants stood barefoot with feet parallel, keeping intermalleolar distance to 10% of the shoulder-to-shoulder distance and with arms free hanging on body sides. Thus, their base of support was relatively wide, about 10 cm, allowing participants to comfortably stand upright. Each block started with a red circular target (diameter of 1.46 degrees of visual angle) projected onto a stationary black background (Figure 1). The target first appeared stationary for 3 sec at the center of the projected screen, which was aligned to the participant’s body midline. This prompted participants to fixate the target, allowing us to ensure that all participants started each trial with the same gaze direction. The target then started moving horizontally, covering a peak-to-peak amplitude of 15, 42, or 95 degrees of visual angle at a constant velocity of 19.9 degrees/sec, and participants were required to accurately visually track this target. The target’s motion signal was a sawtooth wave created in Matlab, with a frequency of 0.48, 0.18, and 0.08 Hz that gave 14.5, 5.5, and 2.5 cycles of horizontal motion in 30 seconds for each target amplitude condition, respectively. After the 30 sec of motion, the target arrived back at the center of the screen, where it remained stationary for another 3 sec, again requiring participants to fixate on it. No instruction about head stabilization was given. Instead, participants were allowed to move their body freely to track the visual target. Each of the three target amplitudes was presented 5 times in a pseudorandom way within each block, resulting in a total of 15 trials per block. Considering that each trial lasted 36 sec, each block took about 10 minutes to complete. The total experiment, including preparations, lasted about 35-40 minutes.

**Figure 1:**
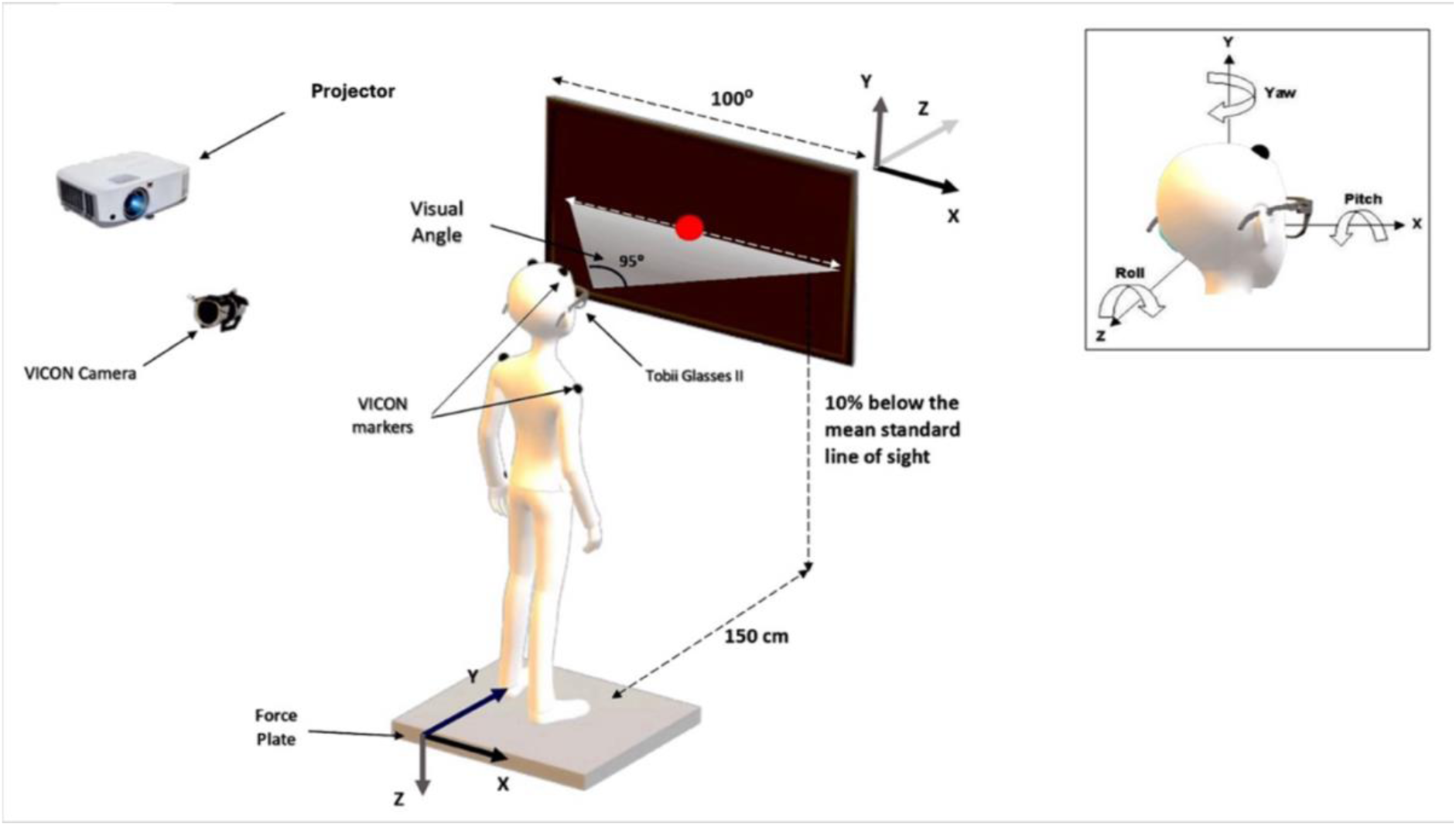
Experimental Set-up. The participant was positioned in front of a wall where the stimulus was projected. The depiction shows the participant standing on a firm surface (i.e. force plate), but they could either be seated on a chair or stand on a piece of foam placed on top of the force plate. The task involved visually tracking a horizontally moving target for 30 seconds. During this period, ground reaction forces, eye movements, as well as head and shoulder kinematics were recorded synchronously.

Experiment 2 was identical except for the details mentioned below. As we were not interested in examining again the influence of the target’s moving amplitude, young and older participants were only exposed to a target that had a peak-to-peak movement amplitude of 42 degrees, equivalent to the mid-range target amplitude of Experiment 1. This was chosen as the most natural condition allowing for flexible behavior since such amplitudes most commonly include the engagement of both the eyes and head for tracking (Hallet, 1986; Daye et al., 2012; Georgiadis et al., 2023). Moreover, participants now performed the task either with the head free to move (as in Experiment 1) or with the instruction to limit their head’s movements when tracking the moving target. More specifically, if a participant employed an average across cycles peak-to-peak head yaw rotation greater than 5 degrees (∼10% of the target’s range of motion), the trial was repeated. This happened in less than 10 trials throughout the complete experiment (< 1% of the total number of trials). The two conditions regarding the head’s engagement, as well as the three postural configurations (seated, firm, foam) were presented in six separate blocks of trials, counterbalanced across participants of each age group. Each block included 5 trials, lasting in total about 3 minutes, for a total experimental time of about 25-30 minutes (including preparations and breaks). Eighteen out of the total 216 trials, were excluded due to occlusions of the head-markers (8.3% of the total dataset), but all participants still had at least three valid trials per condition, and therefore no participant was excluded from the final analyses.

### Data analysis

The mediolateral (ML) and anteroposterior (AP) CoP time series were first computed from the vertical ground reaction force vector and the two moments. The CoP time series were then filtered with a zero-lag 4th-order low-pass Butterworth filter (cut-off frequency of 5 Hz). We calculated the CoP time series only for the two standing conditions because measuring CoP while seated is meaningless. For each trial, we determined the CoP path as the cumulative Euclidean distance covered by the CoP during the 30 sec of pursuing the target, and we then averaged this value across the five repetitions of each of the nine conditions (3 postures x 3 target amplitudes) in experiment 1, and six conditions (2 tasks x 3 postures) in experiment 2.

For the calculation of head rotation about the vertical axis (yaw), the Cartesian 2D linear position coordinates along the lateral and vertical direction of the right and left head markers were smoothened with a zero-lag 4th-order low-pass Butterworth filter (cut-off frequency of 5 Hz) and then converted to angular coordinates using the arctangent function (atan2d) in Matlab R2023b. We then calculated head engagement with respect to the target’s maximal amplitude, allowing us to determine the net effect of the target’s path on head engagement. To this end, we calculated the maximal head yaw rotation for each target cycle and divided this by the actual range of the target’s movement in visual angle. This ratio was then averaged across all cycles within a trial and further averaged across the five trials of each condition to derive the relative head engagement index.

Raw eye-in-head data were extracted from the eye-tracker capturing the participant’s right eye gaze direction and consisting of three-dimensional vector coordinates (x, y, z). Any missing values in the eye data were identified and replaced with interpolated values to maintain data continuity. The interpolation was performed using cubic Hermite interpolation with a modified Akima interpolation (Makima) through the interp1 function in Matlab 2023b. A zero-lag 4th-order low-pass Butterworth filter (cut-off frequency of 5 Hz) was applied to further smooth the gaze data and remove high-frequency noise. Finally, a median filter with a specified filter interval was applied to smooth the data and attenuate blink artifacts. The filter interval parameter was adjusted based on the specific characteristics of the experimental conditions (e.g., target movement amplitude levels) as 30% of the frames within a period of the target’s motion. Rotation angles were then calculated using the arctangent function (atan2d) in Matlab 2023b based on the filtered eye direction vectors.

The overall gaze rotation was calculated as the summation of the head and eye yaw rotations. The spatial coupling between the gaze and the target motion was assessed as the peak of the cross-correlation function between the gaze and the target motion time series. A value of 1 represents perfect spatial coupling between the time series. Overall, given the simplicity and predictability of the visual stimulus, participants demonstrated high cross-correlation coefficient values (above 0.8). In cases where the cross-correlation coefficient was lower than 0.6, data was visually inspected for abnormalities regarding the quality of the kinematic and gaze data. If such trials were deemed to contain corrupted kinematic or gaze data, as this was determined by visual inspection, they were labeled invalid and were excluded from all analyses. This was the case for four participants in Experiment 1, who had at least 15 out of 45 trials excluded, eventually resulting in less than three valid trials in any of the nine conditions. For this reason, we decided to exclude these four participants from all analyses. The remaining 24 participants had at least three valid trials out of the five thirty-second trials within each of the nine conditions. Out of the 1080 valid trials across those 24 participants, 40 were excluded from further analysis (3.7% of the total dataset) due to high variability in the kinematic and/or gaze data rising from calibration issues of the motion tracking system. In these instances, data distortion was distinctively evident, and trials were excluded by visual inspection. No trials were excluded in Experiment 2.

### Statistical analysis

Experiment 1 focused on the effect of postural demands on the head’s engagement in the visual tracking task. Initially, to confirm previous findings that postural sway increases when standing on an unstable than stable surface (e.g., Patel et al., 2008), we compared the CoP path between the foam and firm surfaces with a one-sided t-test. To examine whether the target’s amplitude and the postural demands influence head engagement, we submitted the relative head engagement to a 3 (amplitude) x 3 (posture) repeated measures analysis of variance. We expected a main effect of the target’s amplitude (Freedman, 2008; Daye et al., 2012) with larger head rotations for larger target amplitudes. Our main interest, however, was whether postural demands influence the degree of head engagement during tracking. Main effects of this ANOVA were explored with post-hoc t-tests, corrected with the Holm procedure to account for multiple comparisons. Estimates of effect size are reported using η^2^ for the ANOVA and Cohen’s d for the t-tests. Degrees of freedom were adjusted using Greenhouse-Geisser corrections when the assumption of sphericity was violated. We specified the alpha level to 0.05. All statistical tests were conducted in Jamovi (version 2.5).

Experiment 2 was set to investigate the potential implications of the head’s engagement and postural control in both young and older adults. Firstly, to examine if posture is influenced by aging and whether participants engage their head in tracking, we submitted the CoP path to a mixed 2 (age group: young, old) x 2 (head: fixed, free) repeated measures ANOVA. To examine whether relative head engagement is influenced by aging, possible head restrictions, and postural demands, we conducted a mixed 2 (age) × 2 (head) × 3 (posture) repeated measures ANOVA. Finally, to investigate if visual tracking performance improves when engaging the head, and whether such possible effect depends on aging, we conducted a 2 (age group) x 2 (head task) repeated measures mixed ANOVA on the cross-correlation coefficient between the gaze and target time-series.

## Results

### Experiment 1

Experiment 1 evaluated the effect of the target’s movement amplitude and postural demands on the head engagement of healthy young adults in a visual tracking task. Figure 2 illustrates averaged kinematics of the head, eye, gaze, and target for one participant during the 30 sec of the tracking task across the three different target movement amplitudes (columns) and the three postural conditions (rows). It is evident that gaze (blue curves) highly overlaps with the target’s motion (black lines), suggesting that this participant accurately tracked the target. This was generally the case for the rest of the participants, as reflected in cross-correlations of at least 0.8 between the gaze and the target time series. Furthermore, postural sway was larger when standing on foam than on a rigid surface (*t_71_ = 11.57, p < 0.001, Cohen’s d = 1.36*), confirming that our experiment induced postural challenges in line with previous work (Patel et al., 2008).

**Figure 2:**
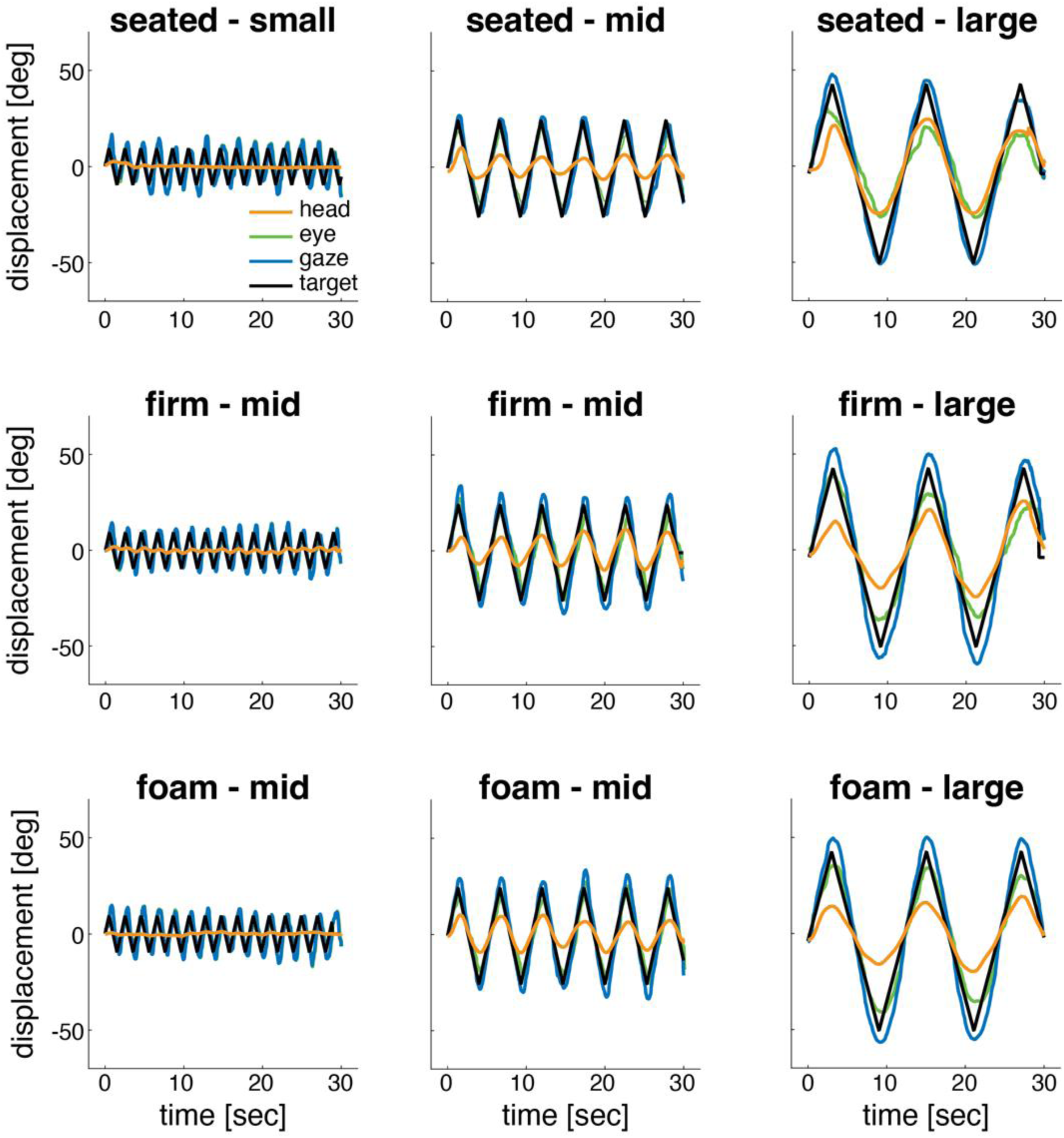
Exemplar kinematics of the head (orange), eye (green), gaze (blue), and target (black) for a single participant across the three postural conditions (rows) and three target movement amplitudes (columns). The data are averaged across trials.

As suggested by Figure 2, and as expected from previous findings (Freedman, 2008; Daye et al., 2012), head yaw rotations during the visual tracking task were larger with wider target amplitudes *(F_1.46, 33.55_ = 39.63, p < 0.001, η^2^ = 0.19*; Figure 3A). Post-hoc pairwise t-test comparisons revealed that head rotations were larger for the large and mid, compared to the small (both *t > 6.44,* both *p < 0.001,* both *Cohen’s d > 1.31*) target amplitudes. However, no significant change in the relative head engagement was observed between the mid and large amplitudes (*t_23_ = 0.93, p = 0.180, Cohen’s d = 0.19*). Importantly, postural demands influenced head engagement during tracking. As shown in Figure 3B, head rotations accounted for about 30-40% of the total target amplitude, and this ratio increased with further postural demands (*F_1.53, 35.12_ = 5.46, p = 0.014, η^2^ = 0.01*; Figure 3). Post-hoc pairwise comparisons showed that head engagement increased from the sitting condition to standing on the firm (*t_23_ = 2.58, p = 0.033, Cohen’s d = 0.52)* and the foam (*t_23_ = 2.83, p = 0.028, Cohen’s d = 0.57*) surfaces. However, although head engagement was descriptively larger when standing on the foam than on the firm surface, this difference was not systematic (*t_23_ = 1.44, p = 0.164, Cohen’s d = 0.29*). No interaction was found between target amplitude and postural condition (*F_2.83, 65.12_* = 0.74, *p* = 0.523, *η^2^* < 0.01).

**Figure 3:**
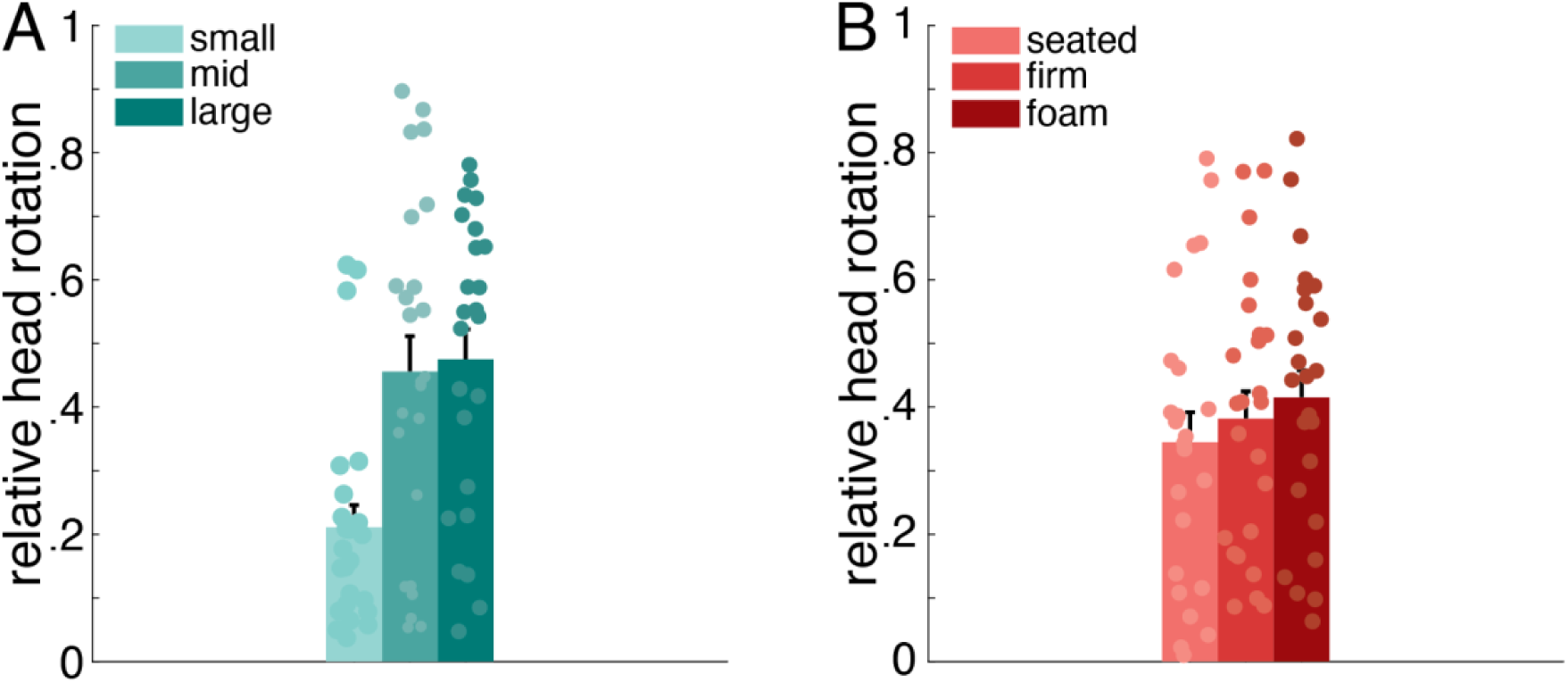
(A) Relative head engagement as a function of (A) target amplitude averaged across the three postural conditions, and (B) postural demands averaged across the three amplitude conditions. Mean values across participants with error bars indicating the standard error.

To further investigate whether head engagement is accompanied by additional body movements, such as trunk rotation, we conducted an exploratory analysis. For this, we determined trunk rotation as the orientation of a line connecting the markers on the acromia. Like the analysis of head engagement, we evaluated trunk rotations relative to the target’s range of motion. Additionally, we calculated the cross-correlation coefficient between the head and trunk time-series to assess the coupling between the rotations of the two segments. Figure 4A shows that postural demands influenced how much participants rotated their trunk (*F_2, 46_ = 28.93, p < 0.001, η^2^ = 0.30*; Figure 4A). Similar to the head engagement results, post-hoc pairwise comparisons revealed that trunk rotation increased from sitting to standing on the firm (*t_23_ = 4.55, p < 0.001, Cohen’s d = 0.93*) and the foam surface (*t_23_ = 8.14, p < 0.001, Cohen’s d = 1.66*), as well as from standing on the firm to standing on the foam surface (*t_23_ = 2.34, p = 0.027, Cohen’s d = 0.48*). The coupling between the head and the trunk also increased as a function of postural demands (*F_2, 46_ = 22.45, p < 0.001, η^2^ = 0.28*; Figure 4B), as it was larger when standing on the firm surface than sitting (*t_23_ = 3.28, p = 0.003, Cohen’s d = 0.67)*, when standing on the foam than when sitting (*t_23_ = 4.63, p < 0.001, Cohen’s d = 0.94),* and when standing on the foam than when standing on the firm surface *(t_23_ = 6.08, p < 0.001, Cohen’s d = 1.24)*.

**Figure 4:**
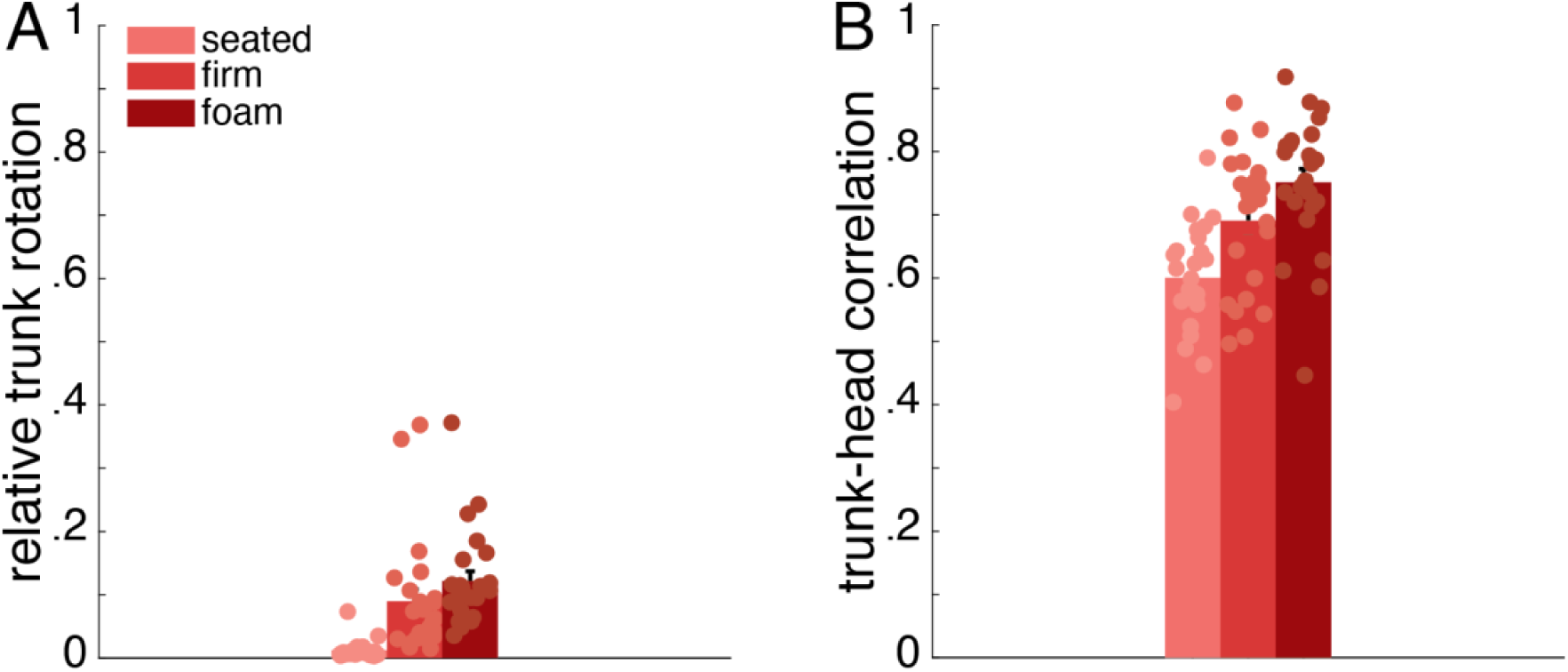
(A) Relative trunk engagement as a function of postural demands, and (B) cross-correlation between trunk and head rotations for each postural configuration. Averages across participants with error bars indicating the standard error. Circles represent individual participants.

We additionally performed an exploratory correlational analysis to investigate the relationship between the amount of head engagement and postural sway. This revealed an overall significant positive correlation between the relative head engagement to the visual task and CoP sway (Supplementary Material; Figure S01). However, this was not the case when looking at each of the experimental conditions (postural demands by target’s movement amplitude) separately (Supplementary Material, Figure S02).

### Experiment 2

Experiment 2 was set to directly assess the interplay between relative head engagement and postural sway. Additionally, we investigated the modulation of age on this relationship. Overall, CoP paths were larger for older than young adults (*F_1, 34_ = 7.51, p = 0.009, η² = 0.06*; Figure 5A) and when standing on the unstable compared to the firm surface (*F_1, 34_ = 5.93, p < 0.001, η² = 0.38*; Figure 5A); both results are in support of previous findings (Thomas et al. 2016; Patel et al., 2008). Moreover, participants’ compliance with the instructions to keep their head still was verified by the increased head engagement in the head-free condition compared to the minimal head rotations during the head-fixed one (*F_1, 34_ = 17.65, p < 0.001, η² = 0.55*; Figure 5B). Finally, participants tracked the moving target with high accuracy in both head-free and head-fixed conditions as reflected in cross-correlations of at least 0.9 between the gaze and the target time series (Figure 5C). These results demonstrate that our experimental manipulations worked as expected.

**Figure 5:**
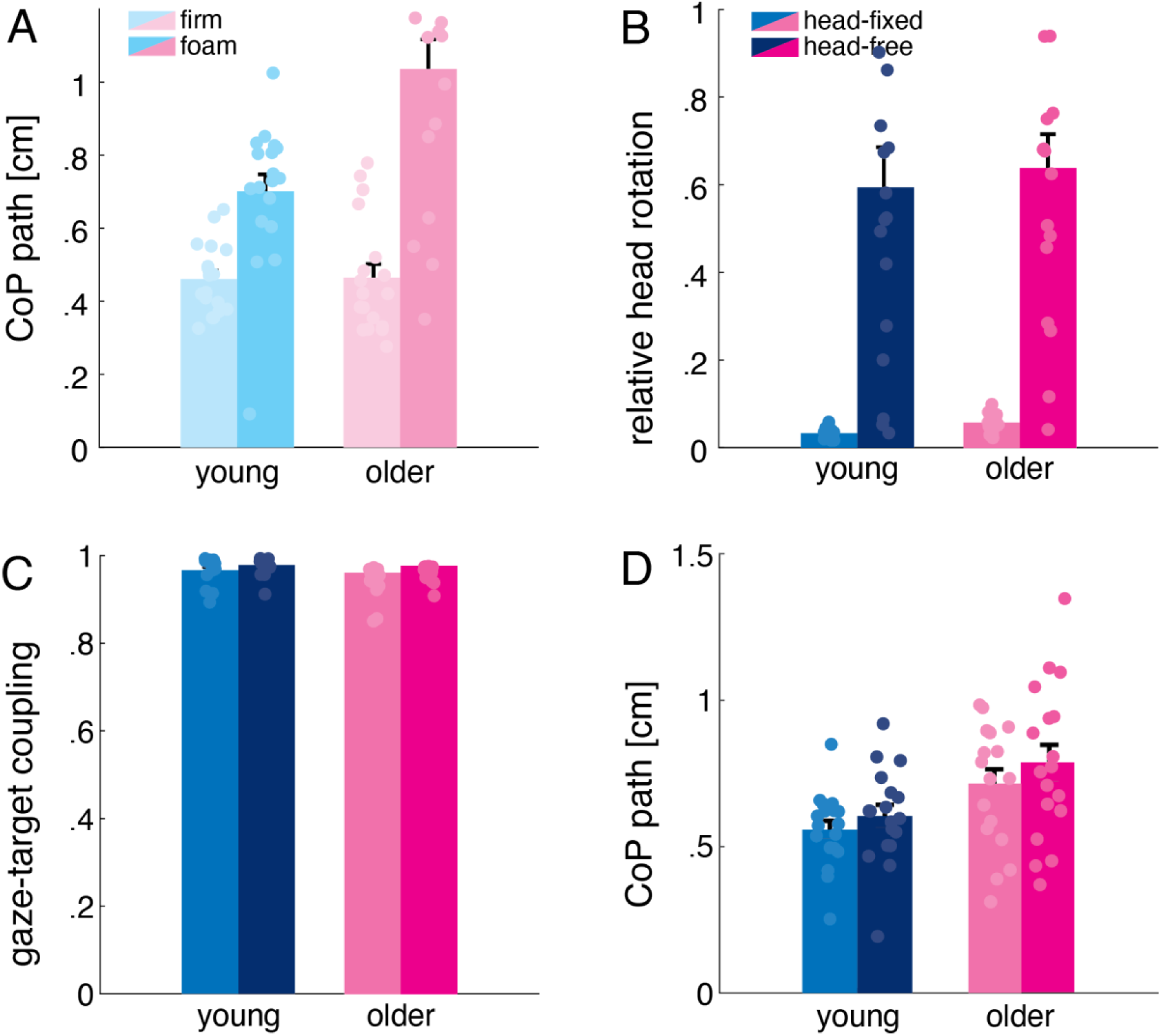
(A) CoP path for each age group and postural configuration, averaged across the two head task conditions. (B) Relative head engagement for each age group and head task, averaged across the three postural configurations. (C) Cross-correlation coefficients for the two tasks and age groups, averaged across the three postural configurations. (D) CoP path for each age group and the two head task conditions, averaged across the three postural configurations. Error bars indicate the standard error of the mean, with small circles representing individual participants.

We first tested whether moving the head per se increases postural sway. Indeed, CoP path was larger when tracking the target with natural head movements than when being instructed to keep the head still (*F_1, 34_ = 6.65, p = 0.014, η² = 0.01*; Figure 5D). There was no effect of age group (*F_1, 34_ = 0.06, p = 0.305, η² = 0.02*) nor an interaction between task and age group (*F_1, 34_ = 0.26, p = 0.610, η² < 0.01*).

Next, we examined whether healthy aging influences the way people engage their head when performing our task. A three-way interaction (*F_2, 68_ = 5.39, p = 0.007, η² = 0.01*; Figure 6) revealed that both age groups slightly increased their head engagement with more demanding postures in the head-fixed condition, but they chose different strategies in the head-free condition: here, young participants increased their head engagement from sitting to standing, as in Experiment 1, while older participants reduced it. This was further verified by a significant posture by age interaction when looking only in the head-free condition (*F_2, 68_ = 3.92, p = 0.024, η² = 0.07)*.

**Figure 6:**
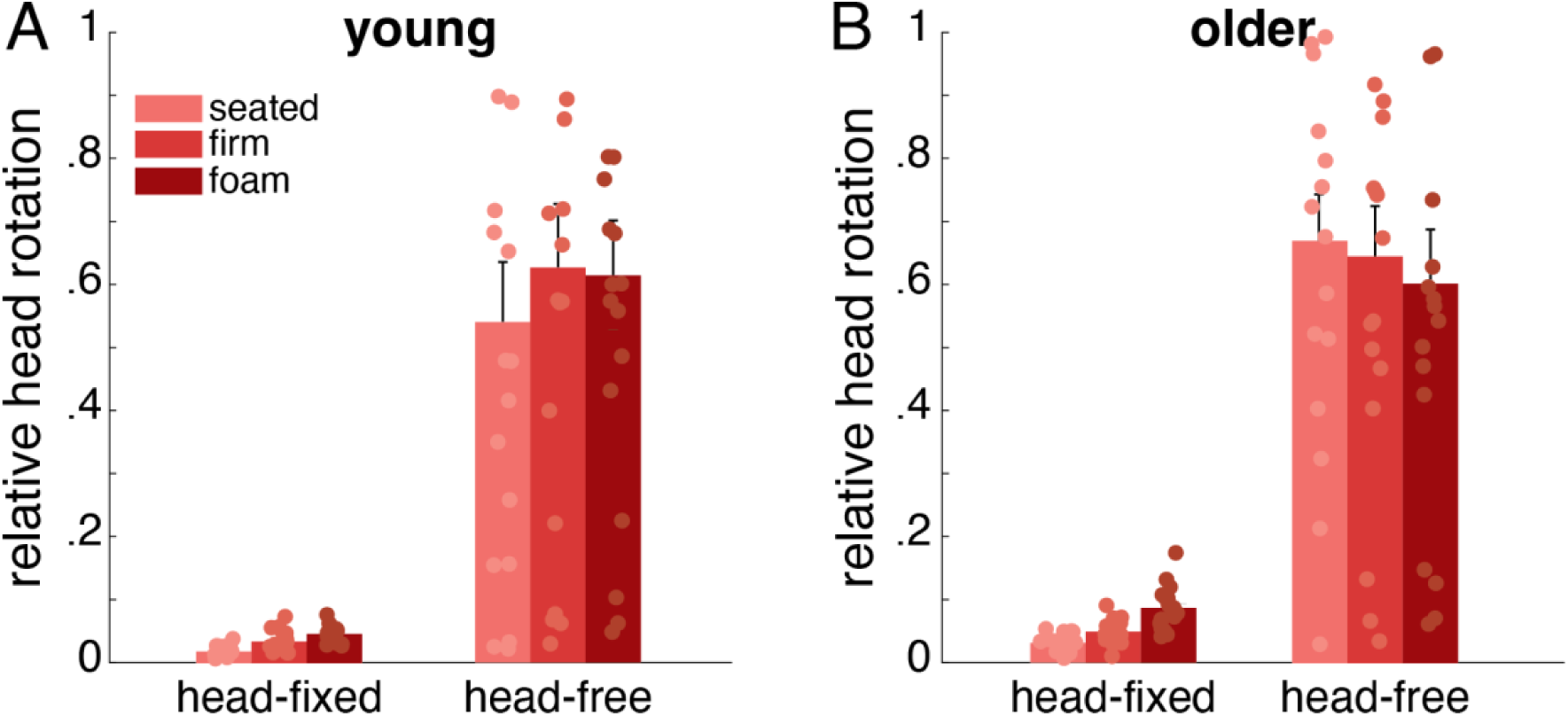
Relative head engagement for the two head tasks and the three postural conditions, separately for (A) younger and (B) older participants. Darker colors indicate more demanding postural conditions (seated, standing, foam). Averages across participants, with error bars indicating the standard error from the mean, and circles representing individual participants.

Finally, we evaluated visual tracking performance through the coupling index between the participants’ gaze and the target’s location across time to account for possible implications of the head’s engagement. Even though all participants tracked the target quite accurately, tracking performance was better in the head-free condition, as the cross-correlation coefficients were higher when allowing than when limiting head movements (main effect of head task: *F_1, 34_ = 7.91, p = 0.008, η² = 0.*05; Figure 5C). There was no effect of age (*F* = 0.23, *p* = 0.628, *η² = 0.03*) nor an age by task interaction (*F* = 0.21, *p* = 0.649, *η² = 0.01)*.

## Discussion

In this study, we investigated the interplay between head engagement and postural control during a visual tracking task. Participants tracked a target moving along three different horizontal amplitudes while they were sitting, standing on a firm, or an unstable (foam) surface. We confirm that participants exhibited larger CoP displacements when standing on an unstable surface, in line with previous work (Patel et al., 2008). We also show that participants engaged their heads more when tracking targets that moved larger amplitudes, supporting previous findings (Freedman, 2008). Our novel finding is that postural requirements increased the degree of head engagement during visual tracking in young adults while they had the opposite effect in healthy older adults.

Experiment 1 revealed that young participants rotated their heads more while visually tracking a moving target in a standing posture compared to when sitting. Although head engagement appeared descriptively stronger when standing on the foam than on the firm surface, the difference was not systematic. This suggests that postural demands alone may not explain the increased head engagement during tracking from a standing posture. One possibility is that participants exploited their motor redundancy by rotating their trunk and head in an en-bloc manner when tracking the target from a standing posture. Indeed, our exploratory analysis showed that trunk rotations increased with postural demands, as also did the coupling between head and trunk movements (cross-correlation of ∼0.7; Figure 4B). However, overall trunk engagement accounted for only about 10% of the target’s amplitude in standing (firm and foam), compared to the head’s contribution of approximately 40% (compared Figures 3B and 4A). Thus, the observed head rotations when standing cannot be fully or solely attributed to trunk rotations. Why, then, did our participants rotate their head more when tracking the target from a standing posture?

Our results indicate that rotating the head for visual tracking might be a beneficial strategy when postural demands arise. It has been shown that head engagement during visual tracking is not only affected by parameters related to the tracking itself, e.g. gaze shift amplitude, but also by sensory and biomechanical factors that prompt the selection of a head orientation where visual information remains centred to the head, e.g., as reflected in bat-and-ball sports (Mann et al., 2013) and steering (Land & Tatler, 2001). A strategy like the latter contributes to keeping the eyes centred at the orbits, which in turn provides flexibility for re-orientating the eyes if necessary and can reduce gaze variability by avoiding extreme eye-in-head orientations. Thus, the additional sensory signals from the head rotations (e.g., neck proprioception) can make the estimates of gaze orientation more reliable (Saglam et al., 2011) and having a more reliable visual representation of the environment can foster postural control (Raffi & Piras, 2019). Hence, we additionally performed an exploratory correlational analysis to investigate the relationship between the amount of head engagement and postural sway. The positive correlation between head engagement and CoP path (Supplementary Material; Figure S01) might stem from larger head movements causing more postural sway. If this were the case, one would expect larger CoP paths for larger target amplitudes because larger target amplitudes were systematically tracked with larger head rotations. However, this does not appear to be the case as there are no systematic relationships between head engagement and CoP path for the separate target amplitudes (Supplementary Material, Figure S02). Instead, we think that the correlation between head engagement and CoP paths is mainly driven by the fact that standing on foam leads to both larger CoP paths and stronger head rotations compared to standing on a rigid surface. It is therefore unclear whether the increase of postural sway with stronger head engagement reflects disturbed postural control or alternatively participants felt free to sway more when having more reliable gaze estimates due to their head rotations. Therefore, we performed a second experiment that directly targeted the relationship between the head’s engagement and the control of posture in two groups with distinct capacities for postural control.

In Experiment 2, we examined the interplay between head movements and postural control in both young and older adults. The results of Experiment 2 demonstrate that visual tracking of a moving target without restricting the head produces greater CoP sways in both young and older adults. Furthermore, we found a significant interaction between age and postural demands. These results support the observed effect of Experiment 1, where more challenging postures led to increased head engagement in young participants. However, older participants did not appear to increase head engagement with more challenging postures, and, if anything, they rather appear to reduce head rotations as postural demands increased. This age-induced modulation could be attributed to the consequences of head rotations on posture, as head rotations impose inertial torques (Fukushima et al., 2008) and downweigh vestibular processing (Roy & Cullen, 2001), both of which can disturb postural control. Young adults, being less susceptible to postural challenges, might have opted to increase their head’s engagement when tracking the target under more demanding postural conditions without much consideration of the effects of head rotation on their posture. In this case, the increased head engagement could reflect a more flexible sensorimotor strategy, where young adults exploit the additional degrees of freedom available in standing to facilitate visual task performance. Indeed, recent literature reports a positive correlation between both CoP sway and performance, as well as head sway and performance, in a visual search task (Bonnet et al., 2024). On the other hand, older adults might have opted to reduce head rotations to prioritize upright stance. Given the reduced capacity of older adults to retain posture and the higher risk of falling, they may have restrained head movement during challenging balance tasks to minimize its destabilizing effects. This strategy could highlight that older adults focus on preserving stability over flexibility or mobility, as has been suggested before (Doumas et al., 2008). This differential effect of aging on head engagement cannot simply arise from an age-related stiffening of the neck because, in the sitting condition, older adults rotated their head as much, if not more, as young participants did (see Figure 6).

In contrast, when participants were instructed to restrain head engagement during visual tracking, both age groups increased head rotation with increasing postural challenge (Figure 6). This spontaneous behavior to engage the head, even when instructed the opposite, could arise as an outcome of the unnatural instruction to keep the head stable. Notably, when no instructions about head movement were provided, both young and older participants employed their head to cover more than half the target’s range of motion across postures (young ∼60%, older ∼65%). Thus, instructing participants to limit head movement during visual tracking could be considered an additional task constraint which imposes a ‘triple-task’ interaction, requiring participants to simultaneously (a) maintain postural control, (b) visually track the moving target, and (c) ensure that they restrict head rotations. The complexity arising from this interaction, as postural demands increase, might have led both young and older participants to regress towards their ‘default’ natural behavior of engaging the head. Therefore, the increase in head engagement with more demanding postures under the head-fixed condition may reflect the reduced ability of both young and older adults to effectively restrain natural head movements in these conditions. Even though this is an assumption, it could explain why older adults increased their head engagement with more challenging postures when instructed to limit head rotations but decreased it when allowed to perform naturally. However, it is important to note that participants’ head rotations, when instructed to restrain their head, did not cover more than 10% of the target’s movement, indicating that these head rotations were minimal (∼4 deg on average) and that the target was mainly tracked by eye movements.

Our results demonstrate that visual tracking with the head fixed leads to smaller CoP sway in both young and older adults compared to head-free visual tracking. This is not another contribution to previous findings showing that postural sway increases during head-fixed smooth pursuit compared to a control fixation condition (Thomas et al., 2016). Our study provides a direct comparison of the same visual tracking task being executed by isolated eye movements and by naturally moving eyes and head together. We show that the head movements employed to facilitate visual tracking are associated with increased postural sway. A larger CoP path is typically associated with poorer static posture (Popović & Popović, 2018) but can also reflect a more flexible postural system with additional degrees of freedom being exploited (e.g., Hatzitaki et al., 2009). Given the dynamic nature of our task—which involves whole-body coordination—and the observed interaction between age and postural demands, the latter explanation is plausible. As previously shown, increased CoP sway during head movements that accompany gaze shifts does not necessarily reflect poorer postural control (Bonnet & Despretz, 2012). An additional factor supporting this possibility is the increased disparity in the proportion of the target’s movement covered by the head between the head-fixed and head-free tasks. If head engagement was detrimental to balance control, one would expect participants that are inherently more sensitive to postural challenges, like older adults, to minimize head engagement to levels like those observed in the head-fixed condition. However, older adults tracked nearly 60% of the target’s movement with their head even in the most demanding postural condition (foam) when free to engage their head, compared to just 0.1% when instructed to keep the head stable. Therefore, even though the increase in CoP sway may not necessarily reflect balance impairments, the two age groups seem to opt for different strategies.

There is evidence that natural head engagement can improve gaze estimates (Saglam et al., 2011) by providing additional input about head orientation. Indeed, our post-hoc analysis showed greater gaze acuity, reflected in the stronger target-gaze coupling in the head-free compared to head-fixed condition for both age groups. This finding provides further insight into the role of head engagement in visual tracking tasks and the origins of the observed differences between young and older adults. In addition, it supports the idea that the two groups differ in how they prioritize competing tasks (Doumas et al., 2008). Since head engagement can enhance gaze accuracy, young adults might benefit from more reliable visual input by increasing head rotation in more demanding postures. In contrast, older adults adopt a suboptimal strategy of reducing head engagement either due to the uncertainty induced by more challenging postures or their diminished ability to manage the mechanical torques associated with head movements.

We conclude that head movements help to visually track a moving target. Although young adults increase their head rotations to track a moving target when postural demands increase, older participants reduce head engagement with greater postural demands. Thus, postural demands seem to differentially affect head engagement in visual tracking in young and older adults, with young adults trading postural sway for head rotations that improve gaze estimates, while older adults reducing head rotations to reduce emergent postural challenges that can compromise upright balance.

## Acknowledgments

We thank Viktoria Wild for assistance with data collection, and Dr. Konstantinos Chatzinikolaou (Aristotle University of Thessaloniki) for the graphical illustration of the experimental set-up (Figure 1). This work was supported by the Deutsche Forschungsgemeinschaft (DFG; German Research Foundation), project number: VO2542/1-1.

## Supplementary Material

Why did participants rotate their head more when tracking the target from a standing than a sitting posture? One possibility is that head rotations provide further sensory signals that assist in visual tracking and improve gaze estimates. As postural control can benefit from reliable visual input, such head rotations could improve posture. Separate correlations between the CoP and the amount of relative head rotations for each of the six conditions did not reveal any systematic relationship (all |r| < 0.508, all p > 0.061; corrected for multiple comparisons; Figure S01), however, when pooling the data across the two standing conditions and the three target movement amplitudes, we observed that participants who rotated their head more also exhibited larger CoP paths (r = 0.31, p < 0.001; Figure 5). A larger CoP path is typically associated with poorer static posture (Popović & Popović, 2018), but can also reflect a more flexible postural system with further degrees of freedom being released (e.g., Hatzitaki et al., 2009).

**Figure S01:**
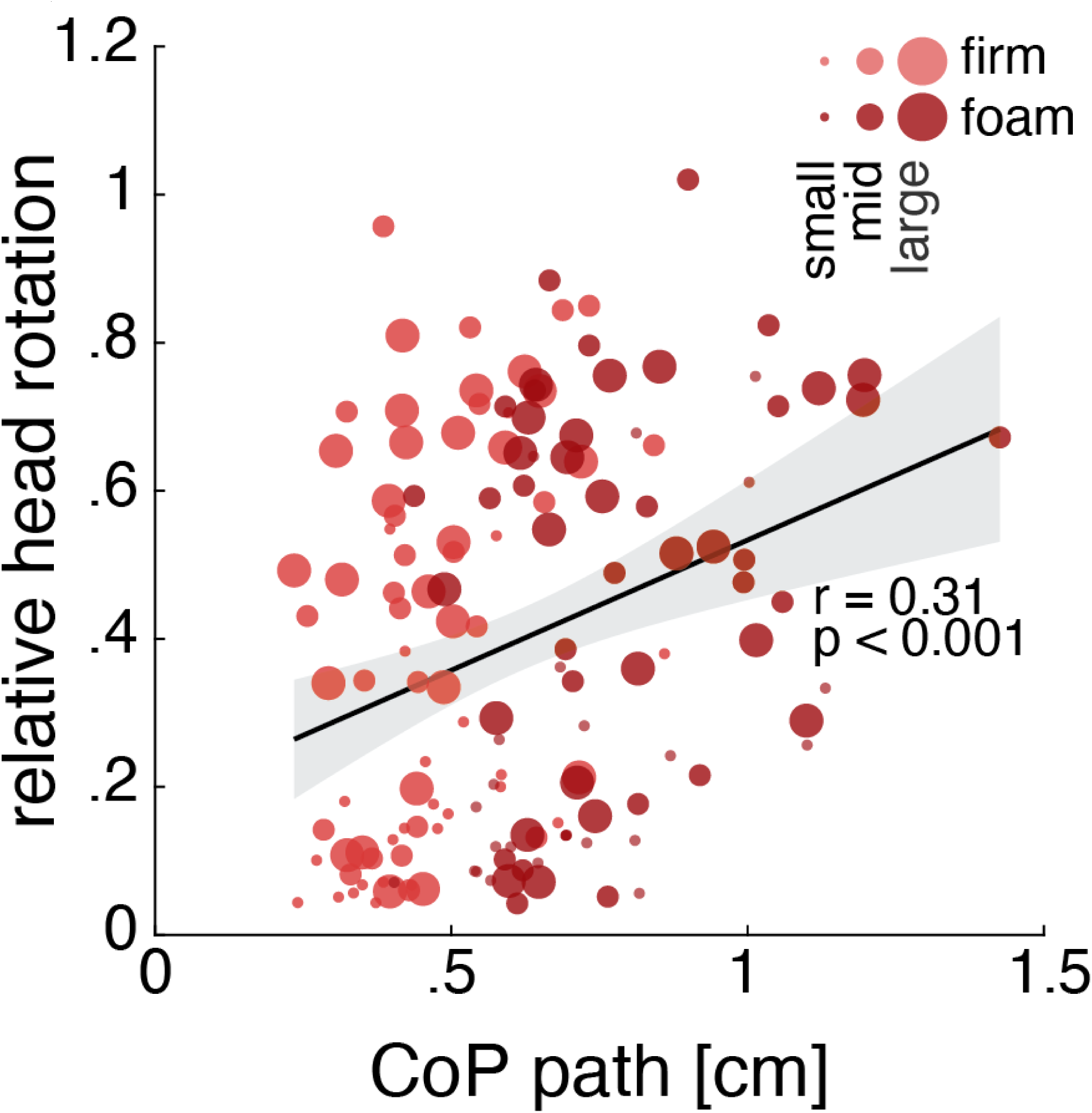
Scatter plot showing the correlation between the CoP path and the relative head engagement, with a fitted regression line (black) and 95% confidence interval (shaded). Data points represent individual values pooled from the two standing postures and the three target movement amplitudes, with the correlation coefficient (r) and p-value (p) displayed within the plot. Postural conditions are color-coded (firm-pink, foam-red), and target movement amplitude is reflected in the size of the scatterdots.

**Figure S02:**
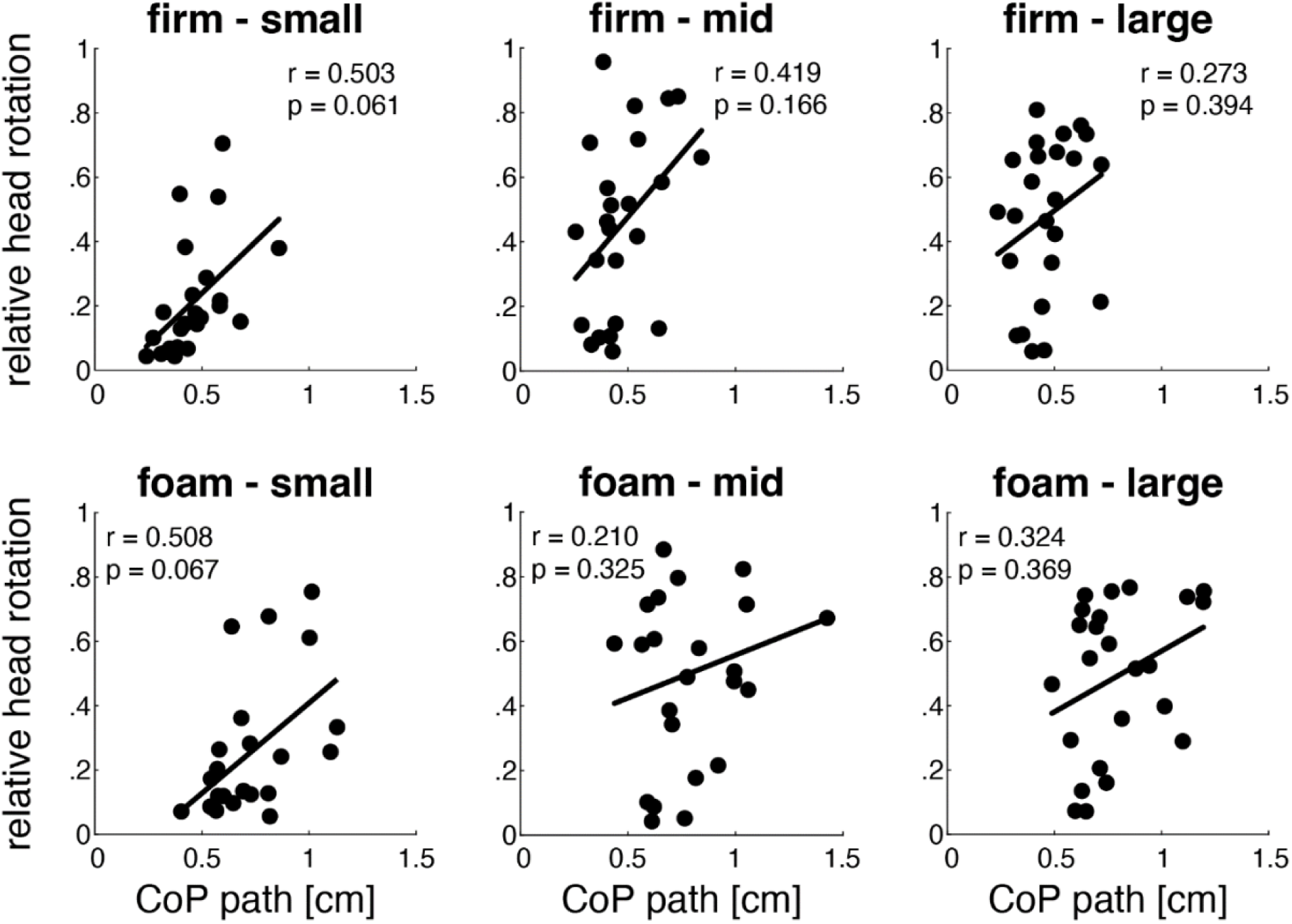
Scatter plot showing the correlation between the CoP path and the relative head engagement, with a fitted regression line (black). Data points represent individual values pooled for each standing posture (firm top row, foam bottom row) and target movement amplitude.

